# Post-reactivation new learning impairs and updates human episodic memory through dissociable processes

**DOI:** 10.1101/320101

**Authors:** Zijian Zhu, Yingying Wang, Jianrong Jia, Liqiang Huang, Yi Rao, Yanhong Wu

## Abstract

Learning of competing information after reactivation has the potential to disrupt memory reconsolidation and thus impair a consolidated memory. Yet this effect has rarely been detected in episodic memory. By introducing an additional retrieving cue to the target memory, the current study detected significant impairment on the reactivated episodic memory, in addition to an integration of new information to the old memory. However, while the integration effect followed the time window of reconsolidation disruption, the impairment effect did not. MEG measurements further revealed alpha power change during reactivation and post-reactivation learning which showed different correlation patterns with the integration and impairment effects, confirming that the two effects relied on different processes. Therefore, post-reactivation new learning disrupts episodic memory but not through reconsolidation disruption. Further findings that the impairment effect was correlated with participants’ voluntary inhibition ability suggest an inhibition-based memory updating process underlying post-reactivation new learning.

## Introduction

Plenty of challenges exist for our memory system. In addition to being stable and accurate, the memory system needs to be dynamic and flexible as the existing information becomes inaccurate, outdated, or even unwanted continuously. In the past decades, researchers have found that a brief reactivation can render a consolidated memory labile again, necessitating a reconsolidation process to restabilize the memory (1). Evidence from three aspects support the role of reactivation. First, amnesic treatments, including pharmacological blockade (2–7) and electroconvulsive shocks (8, 9), impair memory performance when given upon reactivation. Second, extinction training, which diminishes the conditioned response only temporarily, stops the conditioned memory from returning when combined with a brief reactivation (10–16). Third, new competing information learned after reactivation either impairs (17–19) or is integrated into (20–22) the existing memory. Importantly, the effects above show up only after a reconsolidation window closes, suggesting the reconsolidation process being affected.

However, reactivation does not necessarily labialize the consolidated memory. Boundary conditions have been set carefully. For example, Sevenster et al. (23) demonstrates that prediction error during reactivation is necessary for the original memory to become susceptible to interference again. Things are especially complicated in episodic memory (24). In fact, reactivation often have no effect (25) or even improvement (26, 27) on the supposedly labialized memory, suggesting reactivation either does not affect or stabilizes the existing memory. In addition, even if the existing memory becomes more susceptible to interference after reactivation, it does not ensure the effect coming from disruption of the ensuing reconsolidation process. For example, although extinction training upon reactivation stops the conditioning memory from returning in the long run, the conditioning response is diminished even before the reconsolidation process could complete (10). Such effects can be caused by an enhanced version of extinction rather than reconsolidation disruption (28). For episodic memory, a consistent finding that meets the time window criterion of reconsolidation is the memory integration effect. Hupbach et al. (20) found that reactivation rendered the newly learned information more likely to be incorporated into the reactivated memory, an effect that showed up only after the reconsolidation process completed. Such evidence proves that reactivation triggers reconsolidation in the episodic memory. Nevertheless, the reactivated episodic memory was not impaired in Hupbach et al.’s studies (20–22, 29). To sum up, it is still unknown whether reactivation renders episodic memory vulnerable to interference again; and importantly, whether post-reactivation interference disrupts memory reconsolidation needs careful inspection.

To test the effect of reactivation on human episodic memory, we combined the post-reactivation new learning procedure with a double-cue/one-target manipulation (30) and tested both the impairment and integration effects in episodic memory with magnetoencephalography (MEG). Exp. 1 found that post-reactivation new learning had twofold effects. First, the reactivated memory was impaired by the competing information, which was revealed by an independent probe that was not reactivated or interfered with directly. Second, reactivation initiated reconsolidation which allowed the newly learnt information to be integrated into the existing memory. However, while the integration effect showed up only after the reconsolidation window closed (20), the impairment effect was found before reconsolidation could complete. Therefore, reactivation might have triggered another process in addition to reconsolidation which induces memory impairment. This assumption was verified by MEG evidence showing different correlation patterns between the integration and impairment effects with alpha power change during reactivation and post-reactivation new learning. Exps. 2 to 4 further found that the impairment effect could be predicted by participants’ ability to suppress memory retrieval, and relied on intrusion of the reactivated memory. We suggest that reactivation increases intrusion of the original memory, which contradicts with the ongoing new learning and thus triggers involuntary inhibition. The cue-independent memory impairment was caused by inhibition rather than reconsolidation disruption.

## Results

### Memory being impaired by interference upon reactivation

First, we examined whether post-reactivation interference (R-interference) could impair the consolidated memory (Fig. 1A). Participants learned cue-target word associations (e.g. wisdom - plane). After consolidation, the original association was reactivated by retrieving the cue item and then disrupted by learning a cue-substitute (e.g. wisdom - extreme) association. In addition to this classical reconsolidation disruption procedure, two critical manipulations were made. First, the reactivation level was manipulated by controlling the strength of the original memory. Specifically, strong and weak cue-target associations were formed at the beginning, so that presenting the cues individually would induce strong and weak reactivations on the target items respectively. Second, an additional independent cue was introduced. Each target word was paired with two different cue words for learning but only one cue-target series accepted further R-interference training. In the final test, in addition to the trained cue, the untrained cue (i.e. independent cue) was used to retrieve the target item. Thereby, changes on the target item could also be revealed by its independent cue.

**Fig. 1.**
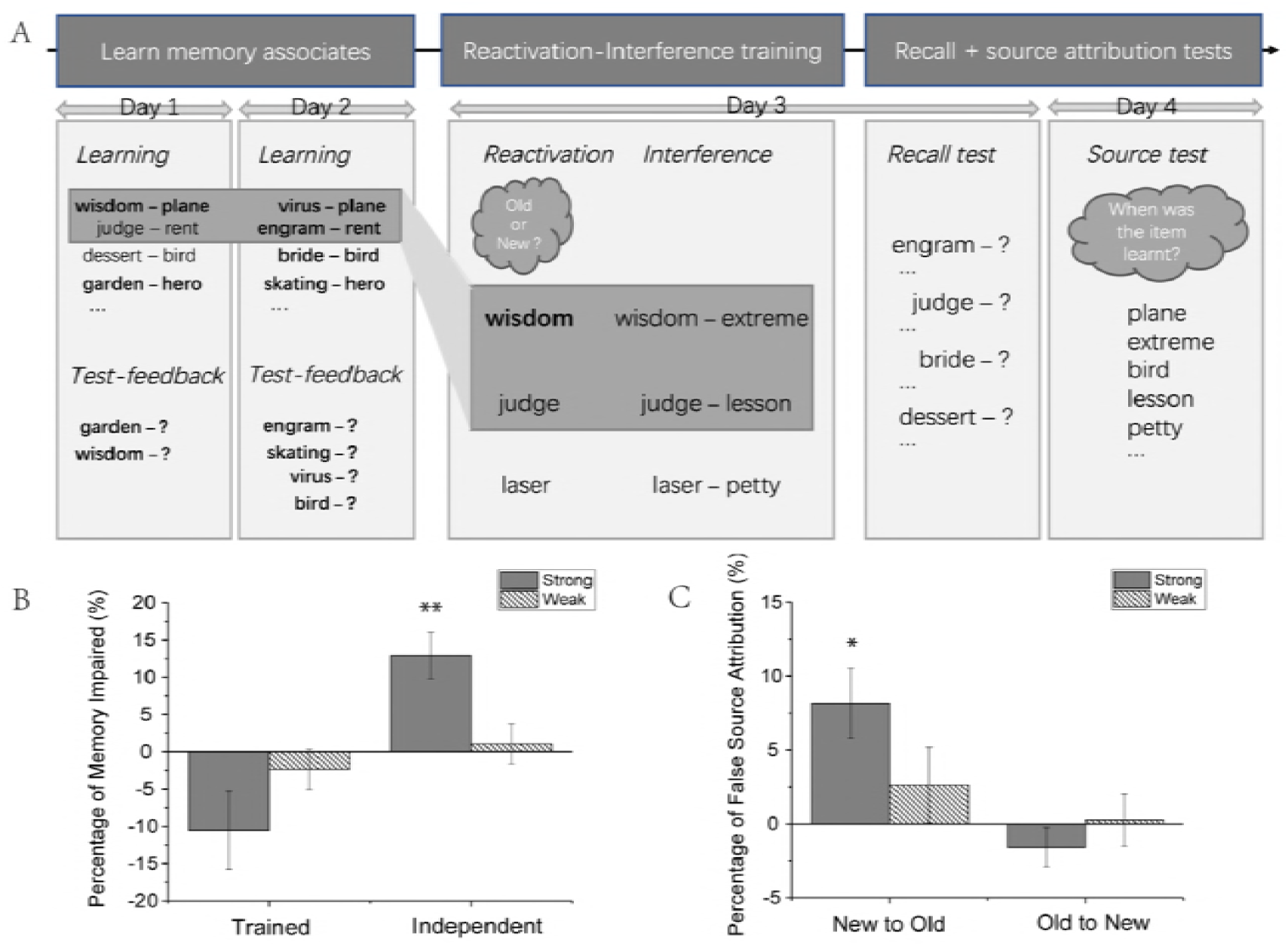
Experimental procedure and behavioural results for Exp. 1. (A) Cue-target word associations (i.e. A-X) were learnt on Day 1. Strong associations (shown in bold) were formed for half of the pairs through a test-feedback procedure to ensure those associations were fully memorized. On Day 2, a second series of cue-target word associations (i.e. B-X) that shared the same targets with those on Day 1 was learnt and fully memorized. On Day 3, a subset of A-X pairs was first reactivated by a recognition test on the cue words and then interfered by pairing the cue words with substitute words for learning. Novel cue words and cue-target associates were included as baseline. A recall test on all the cue words that have been learnt on Day 1 and Day 2 was given right after the reactivation-interference (R-interference) training. On day 4, a source attribution test was given on both the target words and the substitute words, during which participants reported when they have learnt each word. (B) Percentage of memory impaired relative to the baseline condition. R-interference failed to cause memory impairment when tested by trained cues. New learning after strong reactivation caused significant memory impairment when tested by independent cues. (C) Percentage of false source attribution. Substitute words under strong reactivation were misrecognized as the original target words; but the original target words were not misrecognized as substitute words. * *p* < .05, *\*\*p* < .01 (two-tailed t test); error bars, SEM.

We replicated our previous finding (30) that interference under strong reactivation caused significant memory impairment (Fig. 1B, t(18) = 4.11, *p*<.001) when examined by an independent cue that did not accept direct reactivation or interference training. However, no impairment but a trend of improvement were found when the target items were tested by the trained cues (t(18) = −2.01, *p*=.059, marginally significant). The weak reactivation instead failed to affect the original memory, as post-reactivation interference did not cause any memory changes in either the trained- (t(18) = −0.87, *p*=.39) or the independent-cue (t(18) = 0.39, *p*>.70) group. A control experiment with strong reactivation but no interference manipulation was further conducted. No impairment was found any more when tested by the independent (t(19) = 1.05, *p*=.31) or trained (improvement: t(19) = −2.67, *p*=.02) cue. It excluded the possibility that the impairment found in the independent-cue group was caused by reactivation but not interference upon reactivation. Therefore, through introducing an additional dimension, the independent cue, we revealed that reactivation-coupled interference could impair the consolidated episodic memory. However, the impairment effect was found immediately after R-interference training, before the reconsolidation process could complete. Therefore, they were unlikely to be caused by reconsolidation disruption.

### Memory integration through reconsolidation disruption

Incorporating the information learnt during reconsolidation into the original memory has been established as an effect of reconsolidation disruption in human episodic memory (20). In addition to memory impairment, Exp. 1 examined the effect of memory integration caused by post-reactivation interference. A source attribution task was given 24 hours after R-interference training (Fig. 1A), in which both the original and substitute target items were presented and subjects judged when each item was learnt. We calculated the percentage of the interference information (i.e. substitute target items) that was misattributed as the original information (i.e. the original target items). Results (Fig. 1C) showed that more substitutes were misattributed as the original targets in the strong reactivation condition than in the control condition (t(18) = 3.45, *p*=.003). But the original targets were not misrecognized as the interference information (strong reactivation: t(18) = −1.19, *p*=.25; weak reactivation: t(18) = 0.15, *p*=.88). Therefore, consistent with previous findings, reactivation at a certain level renders the newly learnt information to be integrated with the original memory (20–22).

A control experiment was performed to test whether the confusion effect accorded with the time window of the reconsolidation process. Consistent with Hupbach et al.’s (20) control condition, the source attribution test was given immediately after R-interference, before the reconsolidation process could complete. No more confusion was found in the strong reactivation than in the weak reactivation (t(17) = 0.46, *p*=.65) or the control (t(17) = −0.70, *p*=.49) condition. Therefore, introducing interference information upon reactivation leads to the new information being integrated into the reactivated memory through the reconsolidation process.

### Alpha-band activity during reactivation signalling whether to impair or to integrate

Based on the results above, the impairment and integration effects might rely on different processes. Especially, memory integration occurred through the reconsolidation process, while the immediate memory impairment was unlikely to be caused by reconsolidation disruption. To understand what happened during reactivation and interference and to identify the neural activity signalling the impairment and integration effect, we recorded ongoing brain activity using MEG during the R-interference phase. Time-frequency analysis was performed on planar sensors for each trial separately, which was then averaged over trials for each condition and log-transformed.

Six combined planar sensors that showed the largest difference between the strong reactivation and baseline condition at the group level were selected from the posterior occipital area (Fig. 2A). As shown in Fig. 2B, strong reactivation was accompanied with alpha-band (8 – 12 Hz) power decrease when compared with the baseline condition (i.e. seeing novel words). The time of interest was selected between 1.0 and 1.6 s after strong reactivation which showed significant alpha-band power decrease (t(18) = −2.19, *p*=.04). The same trend was also found in the weak reactivation condition when compared with the baseline condition, though not significant (supplementary information, Fig. 2). We further tested the correlation between the neural activity change and the memory change across individuals. We observed a positive correlation between the degree of alpha-band change during reactivation and the degree of memory impairment in the independent-cue group (Fig. 2C, r(17) = 0.638, *p*=.006). In contrast, alpha power change during reactivation was negatively correlated with the level of memory integration through reconsolidation (Fig. 2D, r(17) = −0.490, *p*=.046). Namely, more alpha power decrease predicted more memory integration and less memory impairment. Therefore, the alpha-band activity upon reactivation indicates whether the ensuing interference information is to impair or to be integrated into the original memory. This dissociation suggests the two effects result from different process, which complies with the different time windows of the two effects.

**Fig. 2.**
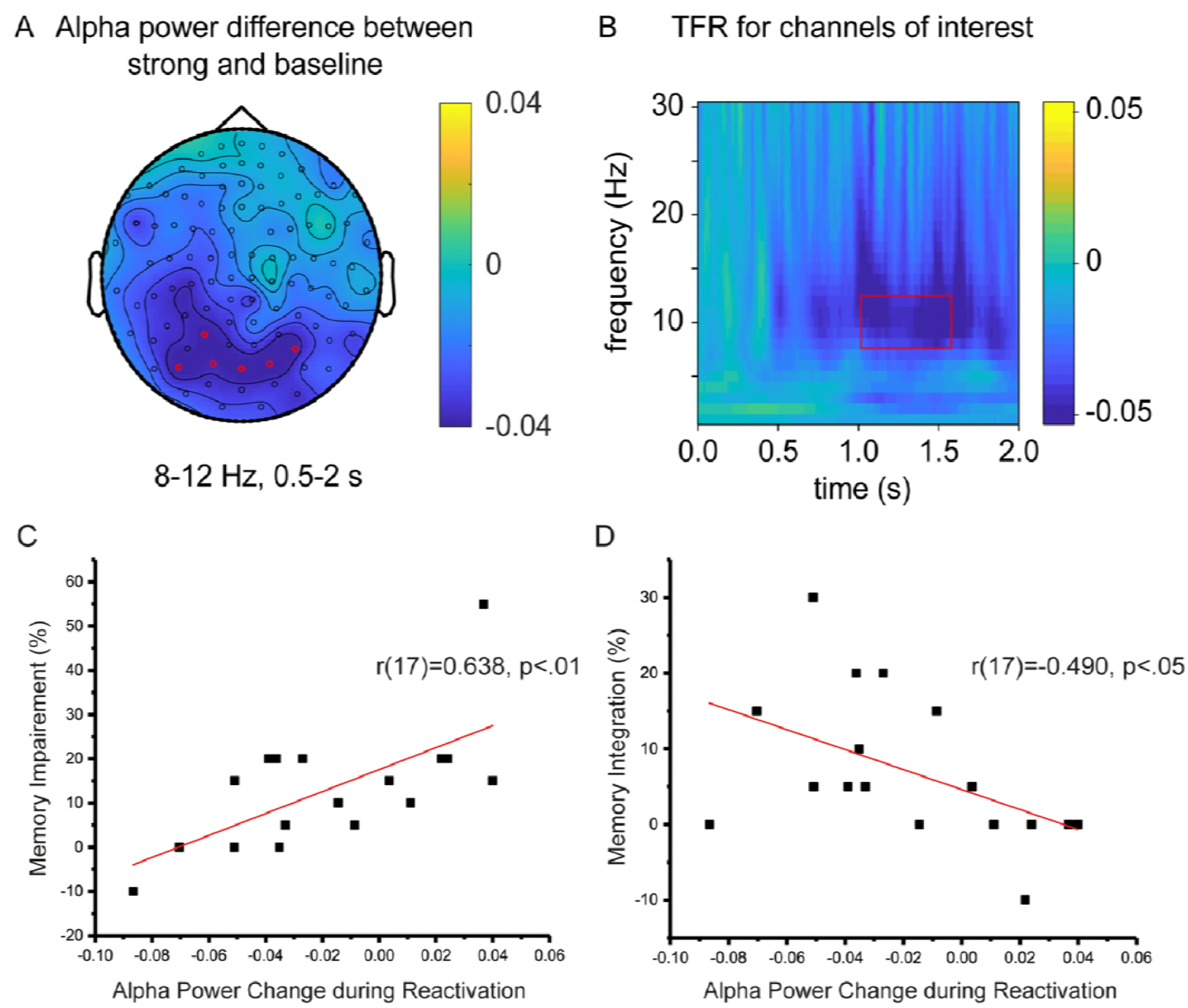
MEG results for the reactivation phase. (A) Grand average (N = 19) topographic map for alpha-band inhibition during reactivation (mean alpha-band (8–12 Hz) power difference between strong and baseline conditions within 0.5 - 2 s). Red circles indicate channels with the strongest alpha decrease. (B) Grand average time-frequency representation (TFR) profile of the selected six channels for the difference between strong and baseline conditions as a function of time (0 - 2s) and frequency (0–30 Hz). Alpha power decreased after reactivation. Red rectangular indicate time of interest (1 – 1.6 s) extracted for correlation analysis. (C and D) Significant correlations were found between alpha power change with the percentage of memory impairment (C) and memory integration (D), but in different directions.

### Alpha- and theta-band activity during interference

Alpha-band activity increased when interference was given. Sensors that showed the largest difference between the strong R-interference and baseline condition were also in the posterior occipital area (Fig. 3A). The time-frequency representation of power revealed stronger 8–12 Hz alpha activity when learning interference information after strong reactivation than learning unrelated novel information. The largest alpha power increase was found at the late interference learning phase, between 3.4 s and 4.0 s (r(18) = 2.05, p = .055, Fig. 3B). The increase showed a trend of negative correlation with the level of memory impairment (r(17) = −0.424, *p*=.09) across participants (Fig. 3C). But no correlation was found between alpha power increase with memory integration (Fig. 3D, r(17) = 0.193, *p*=.46). We also tested the degree of original memory intrusion during interference training. The intrusion degrades gradually over time. We measured the degree of intrusion change across the eight blocks for each subject by calculating its curve slope after linear fitting. Interestingly, correlation (supplementary Fig. 1) was found between the intrusion decrease slope with the impairment of the original memory (r(19) = 0.46, *p*=.05) but not with the integration of new information (r(19) = −0.07, *p*=76).

**Fig. 3.**
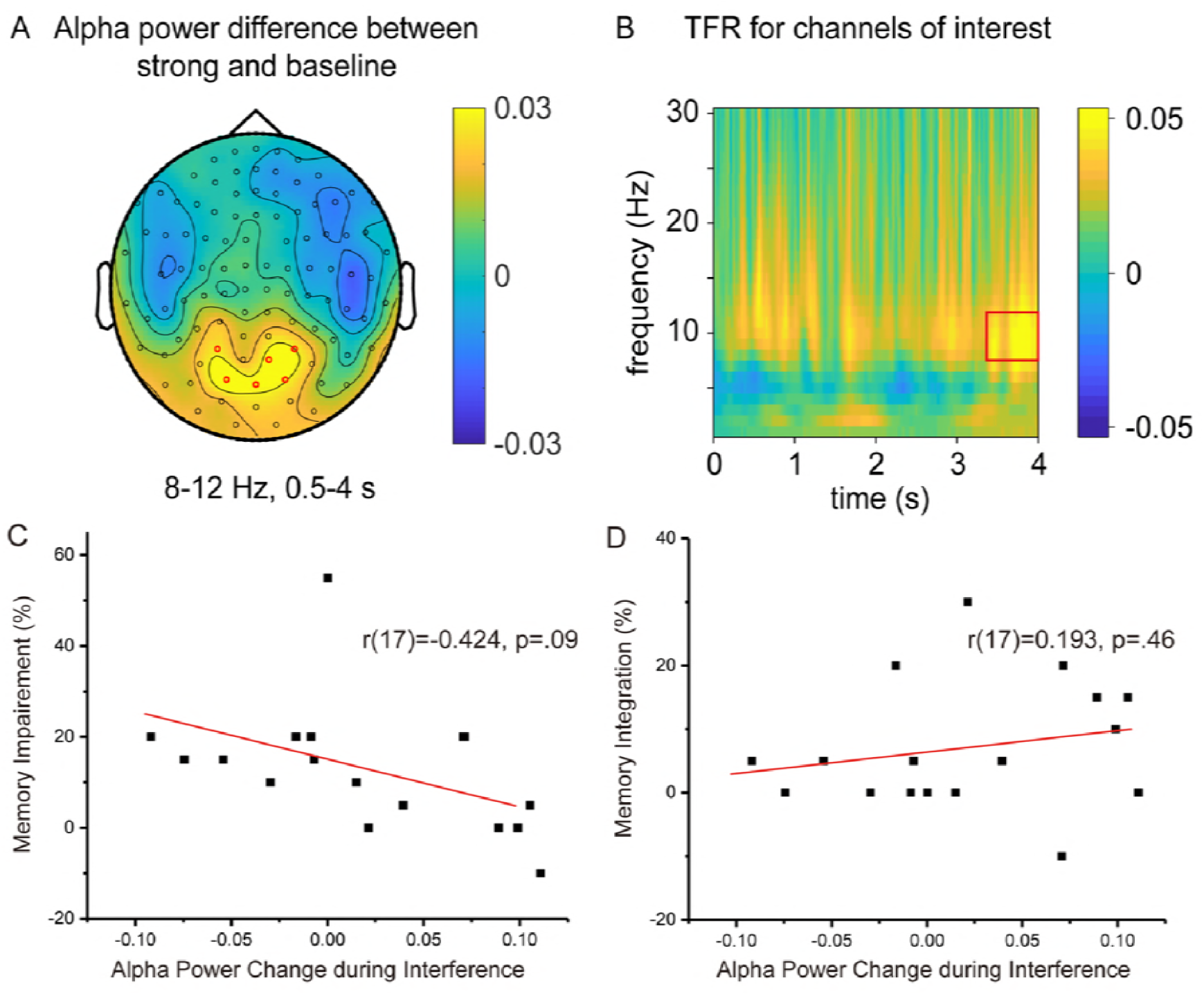
MEG results for the interference phase. (A) Grand average (N = 19) topographic map for alpha-band enhancement during interference (mean alpha-band power difference between strong and baseline conditions within 0.5 - 4 s). Red circles indicate channels with the strongest alpha decrease. (B) Grand average TFR profile of the selected six channels for the difference between strong and baseline conditions as a function of time (0 - 4s) and frequency (0–30 Hz). Alpha power increased and theta power decreased during interference learning. Red rectangular indicates time of interest (3.4 – 4.0 s) extracted for correlation analysis. (C) Marginally significant correlation was found between alpha power increases with the percentage of memory impairment. (D) Alpha power increase during interference was unrelated to memory integration.

Significant changes have also been found at the theta band (4–7 Hz), which is suggested to reflect memory performance (31). As show in Fig. 3B, the theta-band power decreased during interference for strong Reactivation when compared with learning unrelated novel information, complying with the memory impairment found in the strong R-interference condition. However, the theta-band activity change was not correlated with any of the behavioural changes (*p*s>.05).

### Relation of memory impairment by R-interference to retrieval suppression

The memory impairment found in the independent cue by R-interference makes it a promising forgetting approach for real-life events, which are associated with complicated cues. So far, another forgetting approach, retrieval suppression, is known to have similar effect to R-interference (32). Retrieval suppression stops a memory from coming into consciousness by repeatedly suppressing its retrieval. It impairs the target memory through an inhibitory mechanism (33), but its effect has mostly been tested on newly formed memory. To test which approach is likely to be more effective, we compared the two directly on memory that has been consolidated for one week. In Exp. 2, participants learnt double-cue/one-target associations (e.g. wisdom - plane, virus - plane) and accepted interference or retrieval suppression training on a subset of one cue-target series seven days later (Fig. 4A). Before the forgetting training, all of the to be trained cue-target associations were presented to participants to reactivate the existing memory. Memory impairment was found in the independent-cue group by both reactivation-coupled interference (t(30)= 2.70, *p*=.01) and suppression (t(30)= 2.64, *p*=.01) training, when compared with the control condition (Fig. 4C left). More importantly, the degree of forgetting was comparable between the two treatments (t(30)= 0.55, *p*>.05) and significantly correlated with each other across individuals (Fig. 4C right, r(31) = 0.62, *p*<.001). In the trained-cue group, although only retrieval suppression (t(30)= 2.77, *p*=.009) caused memory impairment (Fig. 4B left), the effect was comparable (t(30) = −0.77, *p*=.45) and correlated with each other (Fig. 4B right, r(31) = 0.41, *p*=.02) between the two approaches.

**Fig. 4.**
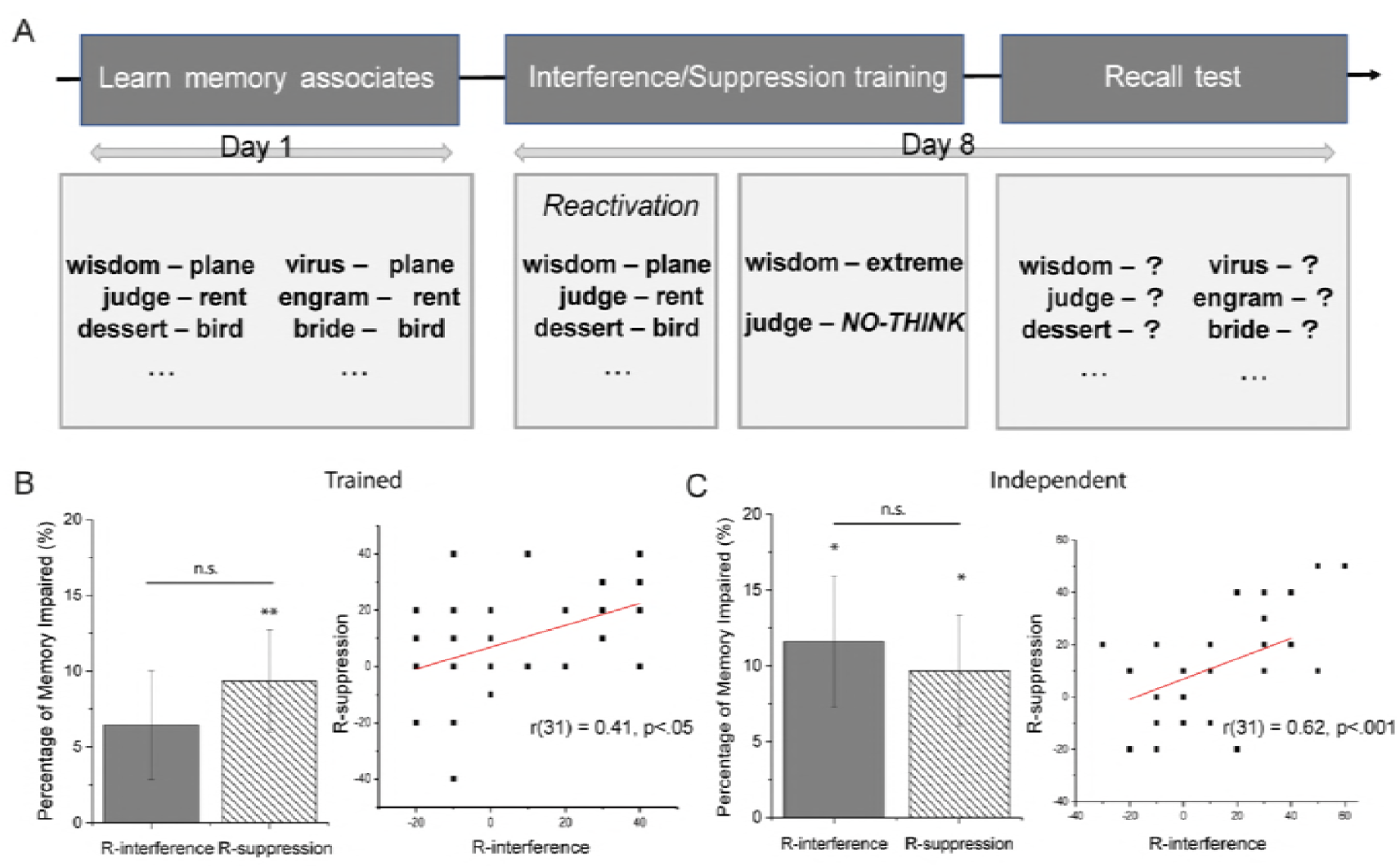
Experimental procedure and results for Exp. 2. (A) Participants learned double-cue/one-target word pairs in the form of A-X/B-X on Day 1. Reactivation was given one week later by presenting all the A-X word pairs for one time. Interference and retrieval suppression trainings were given to a subset of the A-X pairs right after reactivation. Recall test was then given to all the cue words. (B) R-interference and R-suppression caused comparable memory impairment in the trained-cue group (left). Impairment by R-interference was correlated with that by R-suppression (right). (C) R-interference and R-suppression caused comparable memory impairment in the independent-cue group (left). Impairment by R-interference was correlated with that by R-suppression (right). * *p* < .05, ** *p* < .01 (two-tailed t test); error bars, SEM.

### Intrusion but not reactivation is necessary for impairing the consolidated memory

Based on the correlations found above, we hypothesize that the two approaches might share some common mechanism. Considering that retrieval suppression impairs the target memory by suppressing its intrusion (34, 35), the impairment by R-interference might also result from effects induced by intrusion. We speculate that reactivation might increase the original memory intrusion during interference which induces retrieval suppression. In such cases, intrusion rather than reactivation of the original memory causes memory impairment. To test such hypothesis, we performed another two experiments, one (Exp. 3) with and one (Exp. 4) without the reactivation manipulation. Intrusion of the original memory was collected by asking participants to report whether the original target came to their mind during interference/suppression training. Again, with reactivation, impairment was caused by both R-interference (t(33) = −3.23, *p* = .003) and R-suppression (t(33) = −3.29, *p* = .002) in the independent-cue group (Fig. 5A), and the two effects were correlated with each other across participants (Fig. 5C, r(34) = 0.63, *p*<.001). Only R-interference caused memory impairment in the trained-cue group (t(33) = −2.64, *p* = .013), and its effect was still correlated with the impairment by R-suppression (t(33) = −1.41, *p* = .26) in the trained-cue group (Fig. 5B, r(34) = 0.58, *p*<.001). However, when the consolidated memory was not reactivated (Exp. 4), interference (i.e. NR-interference) did not cause any memory impairment in either the trained- (t(32) = −0.81, p = .43) or the independent-cue (t(32) = −1.26, p = .22) group (Fig. 5D). Retrieval suppression (i.e. NR-suppression) managed to induce impairment to the independent-cue group (t(32) = - 4.45, *p*<.001), though no effect was found in the trained-cue group (t(32) = 0.13, *p*=.90). Therefore, well consolidated memory is resistant to interference, but is still vulnerable to retrieval suppression.

**Fig. 5.**
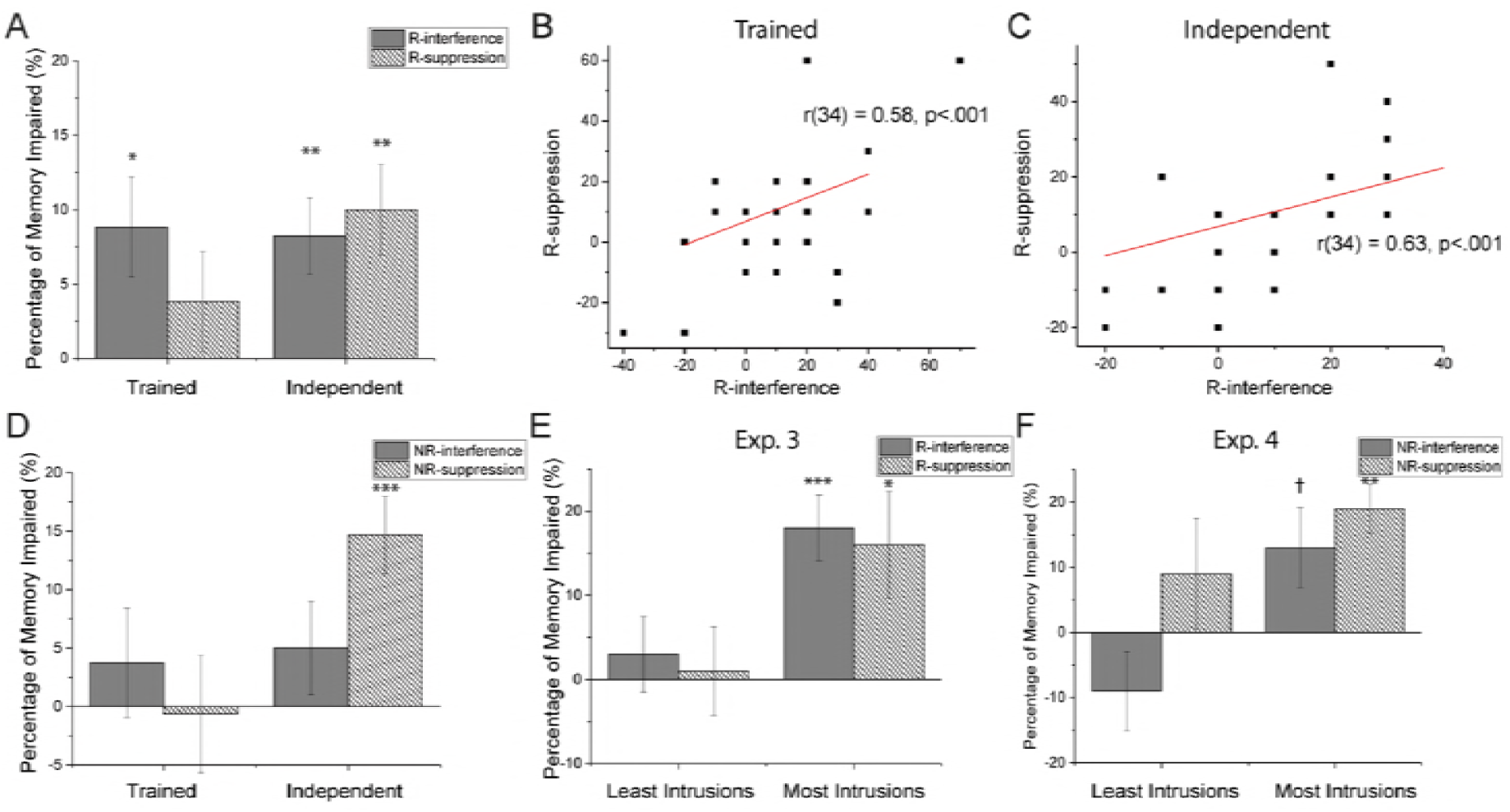
Results for Exp. 3 and Exp. 4. (A) R-interference caused memory impairment in both the trained- and the independent-cue groups. R-suppression caused memory impairment in the independent-cue group. (B) Percentage of memory impaired by R-interference was correlated with that by R-suppression in the trained-cue group. (C) Impairment by R-interference was correlated with that by R-suppression in the independent-cue group. (D) Interference without reactivation (NR-interference) failed to induce memory impairment in either the trained- or the independent-cue group. Retrieval suppression (NR-suppression) managed to impair the consolidated memory in the independent-cue group. (E) Both R-interference and R-suppression failed to cause memory impairment for participants (N = 10) who experienced few original memory intrusions. R-interference and R-suppression caused memory impairment for participants (N = 10) who experienced original memory intrusions a lot. (F) NR-interference and NR-suppression failed to cause memory impairment for participants (N = 10) who experienced few original memory intrusions. NR-interference and NR-suppression caused at least marginally significant memory impairment for participants (N = 10) who experienced original memory intrusions a lot. † *p* < 06, * *p* < .05, *** *p* < .001 (two-tailed t test); error bars, SEM.

Next, we tested whether intrusion was the direct cause for memory impairment. To test the effect of reactivation on intrusion, we calculated the percentage of memory intrusion in the first run, which took place right after reactivation (if there was), of the interference/suppression training. Less intrusion happened when without reactivation, but only during interference training (t(62) = 3.37, *p*=.001; for suppression training: t(62) = 0.37, *p*=.72). We then extracted participants who experienced the least and most intrusion of the existing memory during interference/suppression training, ten for each group. We found that, in Exp. 3, even with explicit reactivations, the ten people who had the least intrusion failed to forget the original targets under R-interference (t(9) = −0.67, *p*=.52) or R-suppression (t(9) = −0.19, *p*=.85). Impairment was found in people who had the most intrusions in Exp. 3 (R-interference: t(9) = 4.63, *p*<.001; R-suppression: t(9) = 2.52, *p*=.03). Likewise, in Exp. 4, participants who experienced the least intrusion had no forgetting effect under either NR-interference (t(9) = −1.49, *p*>.05) or NR-suppression (t(9) = 1.06, *p*>.05). In vast contrast, although without reactivation (Exp. 4), participants who experienced the most intrusion had at least marginally significant forgetting effect under both interference (t(9) = 2.11, *p*=.06) and retrieval suppression (t(9) = 5.02, *p*<.01) training. Consistent with this, the chance of memory intrusion was positively correlated with the degree of memory impairment by both R-interference (supplementary Fig.3, Exp. 3: r(32) = 0.37, *p*=.04) and NR-interference (supplementary Fig.4, Exp. 4: r(32) = 0.43, *p*=.02). Therefore, the intrusion of, rather than the explicit reactivation on, the original memory is essential for interference to impair the consolidated memory.

The correlated forgetting effects and the common reliance on original memory intrusion suggest the two approaches might share some common mechanisms. We suggest that a suppression process is likely to underlie post-reactivation new learning. In the context of interference learning, intrusion of the original memory (partly due to reactivation) contradicts with the ongoing task; to overcome this contradiction, voluntary suppression on the intruding memory is initiated automatically.

## Discussions

Tremendous changes can be made to consolidated memory with the benefit of reactivation. Competing information learnt upon reactivation impairs the existing memory and at the same time updates the memory by incorporating newly learnt information. Our results demonstrate that these two effects rely on distinct processes. MEG recording reveals alpha-band activity decrease during reactivation that is correlated with the two behavioural effects, but in opposite directions. Alpha-band activity increases during interference which reflects the degree of original memory impairment. Meanwhile, while the integration effect shows up only after the reactivated memory is reconsolidated, the impairment effect is found before the reconsolidation process completes, suggesting only the former effect being caused by disruption of memory reconsolidation. The potential mechanisms underlying the impairment effect was then investigated. We compared the impairment by R-interference with that by retrieval suppression, and found cue-independent impairments that are correlated with each other under the two manipulations. In addition, impairment by both R-interference and retrieval suppression relies on memory intrusion rather than reactivation. We thus suggest that retrieval suppression might underlie the impairment effect induced by R-interference. To conclude, reactivation not only initiates a reconsolidation process but also increases intrusion of the existing memory which might trigger involuntary inhibition and impair the reactivated memory.

The current study suggests that reactivation not only induces reconsolidation. The effect by post-reactivation intervention follows a strict time window which serves as evidence for the existence of a reconsolidation process. Any memory change, should it be caused by reconsolidation disruption, follows this time window (1). Consequently, the effect of reconsolidation disruption appears only after the reconsolidation process completes, which takes at least several hours (a recent study further suggested the need of sleep to complete the reconsolidation process (36)). This time window criterion was followed by pharmacological studies, but not exactly in studies using behavioural interference. In conditioning memory, although not different from the effect by extinction without reactivation, extinction after reactivation eliminated conditioning response in immediate test before reconsolidation completes (10). With the assistance of an independent cue, the current study found an immediate forgetting effect by R-interference in episodic memory which did not exist in NR-interference (30). Therefore, the effect violates the time window limit of reconsolidation disruption. This is consistent with Chan and LaPaglia (2013)’s finding that reconsolidation disruption procedure impaired episodic memory also before reconsolidation process could complete (19). In addition, the current study demonstrated different neural signals for the impairment effect and the integration effect, a canonical effect that has been verified to be caused by reconsolidation disruption (20). Therefore, reactivation is likely to induce an additional memory modification process that is independent of reconsolidation. This finding suggests that memory impairment found in R-interference in remote test may not only result from reconsolidation disruption. Time window should be carefully tested before attributing any effect to reconsolidation disruption.

So far, it is well known that reactivation labialize a consolidated memory because interference upon reactivation causes more memory impairment, but it is unknown what underlies this labialization. Our results offer some possibilities for inhibition underlying this process. First, we demonstrated that retrieval suppression, an inhibition–based forgetting approach, works on memory that has been consolidated for one week, which provides feasibility for using inhibition to explain the impairment found in R-interference. Second, common characteristics exist between the two approaches. Retrieval suppression impairs the target memory by repetitively suppressing its retrieval. Its forgetting effect is thus dependent on the intrusion of the original memory (34, 35). In R-interference, reactivation increased the chance of original memory intrusion during interference, which predicted the degree of memory impairment. More than that, a characteristic that discriminates retrieval suppression with traditional interference approaches is that its effect is not restricted to the trained cue (32, 37). Impairment in the independent cue is also found in R-interference in our study. Cue-independent impairment has been used as evidence for the target memory trace being impaired (32), which further explains the reason why R-interference stops the reactivated memory from returning in the long run (12). Forcato et al. (2007) examined the strength of the original memory after R-interference (38). They found that retrieval on the original memory did not cause retrieval-induced forgetting any more, indicating attenuation of the original memory trace. We therefore speculate that inhibition happens in R-interference, which causes memory impairment in immediate test. However, our results suggest existence of inhibition in R-interference but do not exclude the possibility that the impairment also comes from reconsolidation disruption or the combination of reconsolidation disruption and inhibition.

To our knowledge, very few studies have measured brain oscillations under reactivation-coupled interference in episodic memory. Critical results were found at the alpha band during both reactivation and interference. Alpha-band activity is known to respond to a stimulus or task demands either with a decrease or increase in power (39). The most general observation is that task-related brain regions exhibit alpha power decrease, whereas regions associated with task irrelevant and potentially interfering processes exhibit alpha power increase (40, 41). Alpha power increase is thus suggested to reflect inhibition. In the current study, alpha power decreased significantly during reactivation, which is consistent with the previous findings that alpha power decreased for successful encoding and retrieval of long-term memories (42, 43). And successful reactivation of the original memory was found to cause integration of newly learnt information (29). This corresponds with our finding that alpha power decrease was correlated with memory integration through reconsolidation. Future studies could further examine the relationship between alpha power decrease and memory reconsolidation. In contrast, alpha activity increased during interference learning which was correlated with the degree of original memory impairment (Fig. 3C). Moreover, alpha power increase was also correlated with the amount of original memory intrusion (r(17) = 0.587, *p*<.05) during interference learning, which, as the current study suggested, triggered inhibition and caused forgetting. We suggest that the alpha power increase during interference learning might reflect the maintenance of intrusion information. This is consistent with the findings that alpha-band activity indicates the number of working memory units in maintenance (44, 45). The current study also inspected decrease of theta power during interference. Theta-band activity has been viewed as a signature for memory encoding and formation (41). The decrease of theta power accorded with the memory impairment effect. Yet theta power decrease was not correlated with the degree of memory impairment, which might be due to new information learning happened at the same time.

In addition to the possible mechanisms underlying reactivation-coupled interference, the current study establishes that retrieval suppression is effective on well-consolidated memory. While it is known that reconsolidation disruption only applies to consolidated memory, retrieval suppression has been tested mostly on newly-formed memory. A recent study reported not finding impairment by retrieval suppression in consolidated memory. Combining with the fMRI results, Liu et al. (46) suggest that consolidation promotes the assimilation of newly acquired memories into more distributed neocortical regions, making memories more resistant to suppression through prefrontal-hippocampal inhibitory pathways. Our test on memory that has been consolidated for one week replicated their findings in the trained-cue group (Exp. 4, Fig. 5D). Nevertheless, memory impairment was consistently found when tested with an independent cue (Exps. 2–4). Therefore, we suggest that retrieval suppression works on well-consolidated memory. This is also consistent with the fact that retrieval suppression is effective in impairing autobiographical memory (47–49).

The current study observes both integration of new information and impairment on the original memory when interference information is given upon reactivation. However, taking advantage of a double-cue/one-target technique, we demonstrate that the impairment effect appears even before the reconsolidation process completes. This finding call attention to the impairment effect by post-reactivation new learning in episodic memory studies. Considering the differences between episodic memory and other memory formats, it is yet unknown whether this effect applies to conditioning memory, procedural memory, and so on.

## Materials and Methods

Nineteen participants (aged 19–31 years, 12 females) participated in the MEG experiment (Exp. 1), with 20 and 18 participants being further recruited for the two control experiments. Thirty-two (aged 19–31 years, 24 females), 34 (aged 19–28 years, 21 females) and 32 participants (aged 19–28 years, 21 females) participated Exps. 2, 3 and 4, respectively. They were all native Chinese speakers, with normal reading and comprehension ability, and no known neurological disorders. Participants gave written, informed consent in accordance with the procedures and protocols approved by the human participant review committee of Peking University. Detailed methods are provided in SI Materials and Methods.

## Supplementary Methods

### Exp. 1

#### Participants

Nineteen participants (aged 19–31 years, 12 females) were recruited from Peking University, Beijing, China. They were all native Chinese speakers, with normal reading and comprehension ability, and no known neurological disorders. The sample size was determined by a priori power analysis.

Participants gave written, informed consent in accordance with the procedures and protocols approved by the human participant review committee of Peking University.

#### Stimuli

Stimuli were 160 Chinese cue-target word pairs (e.g. wisdom-plane). Each target word was paired with two different cue words (e.g. wisdom-plane and virus-plane), thus two series of word pairs, composed of 80 word pairs each, existed in the form of A-X and B-X. Both A-X and B-X pairs were learnt, but only the A-X pairs accepted reactivation and interference training. The 80 A-X pairs were divided into four groups, 20 pairs each: strong reactivation, weak reactivation, baseline for strong reactivation, and baseline for weak reactivation. The 80 B-X pairs were divided into the same four groups accordingly.

Forty novel words were used as substitutes for post-reactivation new learning, and were paired with cue As in the strong and weak reactivation groups to interfere with the original association (e.g. wisdom-extreme). Another 20 cue-target filler pairs were included during interference learning as a control condition. All the words were frequency-balanced two-character Chinese words.

#### Procedure

The experiment consisted of three phases: associative learning, post-reactivation interference (R-interference), and testing.

##### Associative learning

Participants learned 160 cue-target word pairs (80 A-X and 80 B-X pairs). The A-X pairs were learnt at two strength levels: strong and week. For the strong memory group, 40 A-X pairs were learnt through a learning and test-feedback session until all the right-hand target words could be recollected correctly when given the left-hand cue words (see (Zhu et al., 2016). For the weak memory group, 40 A-X pairs were presented to the participants for four times, 4 s each time. Different from A-X pairs, all B-X pairs were learnt through the same learning and test-feedback session as the strong A-X pairs, so that all of the B-X pairs were fully memorized. In case the different memory strengthens of the A-X associations affected B-X pairs, controls for different conditions were prepared separately. That is, the 40 strong/weak A-X/B-X pairs were split into two groups, one accepted further R-interference training and the other one did not go through R-interference and served as control for the first one.

Learning was completed in two days. All the participants learnt A-X pairs on Day 1 and B-X pairs 24 hours later. This manipulation ensured that the memory strengths for B-X pairs were as equal as possible between the strong and weak groups.

##### Post-reactivation interference

R-interference was conducted in the MEG scanner, 24 hours after learning was completed. Forty A-X pairs (20 strong and 20 weak pairs) along with 20 fillers underwent the R-interference procedure. The remaining 40 A-X pairs (20 strong and 20 weak pairs) did not accept reactivation or interference training and served as control for the corresponding memory strength condition. Reactivation was implemented by a retrieval test. In each trial, a cue A word or a filler word was presented on the screen for 2 s. Participants were asked to judge whether they had seen the word during the initial learning phase internally to avoid influencing the MEG signals. No explicit response was required. Interference was given immediately after reactivation, in which a novel substitute word appeared on the right side of the cue/filler word for 4 s. Participants were asked to memorize the cue-substitute association. Afterwards, subjects reported whether memory for the original target intruded during interference learning on a 4-point Likert scale.

The R-interference procedure repeated eight times, in eight separate blocks. Stimuli for different conditions were presented in a pseudorandom order. No more than three words in the same condition were presented consecutively.

##### Testing

A recall test was given to each participant right after R-interference training. One hundred and sixty cue words were shown on the screen sequentially. Participants tried to recall the target word that was paired with the cue word during the learning phase. A memory source test was given 24 hours later. Target words that have been learnt in the associative learning and R-interference training phases, namely target words from the original word pairs and substitute words from the interference word pairs, were presented to participants. Participants judged when each word was learnt, by selecting from the following choices: *Day 1, Day 2, Day 3*, and *both Day 1 and Day 2*. No time limits were given during either the recall or the memory source test.

##### MEG Recording

Ongoing brain activity was recorded (sampled at 1000 Hz) using a whole-head MEG Neuromag (VectorView™, Elekta Neuromag Oy, Helsinki, Finland) acquisition system. It consists of 306 sensors arranged in triplets of two planar gradiometers and one magnetometer. Before the recordings, four head position indicator coils attached to the scalp determined the head position with respect to the sensor array. The location of the coils was digitized with respect to three anatomical landmarks (nasion and preauricular points) with a 3D digitizer (Polhemus Isotrak system). A custom-made chin set was used to fix the head. The head position with respect to the device origin was acquired before each block and monitored throughout each recording block to ensure that head movements did not exceed 0.5 cm at any time. The temporal extension of signal-space separation (tSSS) method was applied during the pre-processing stage of analysis using Elekta Neuromag MaxFilter software to reduce noise from the external environment (Taulu & Simola, 2006). Two bad channels were manually detected (noisy, saturated, or with SQUID jumps) and excluded for further analysis.

##### MEG Data Analysis

Data analysis was performed in Matlab 2016a (MathWorks, Natick, MA) using the Fieldtrip open source Matlab toolbox developed at the Donders Institute for Brain, Cognition and Behavior, Center for Cognitive Neuroimaging, Nijmegen, The Netherlands (Oostenveld, Fries, Maris, & Schoffelen, 2011) (http://fieldtrip.fcdonders.nl), and custom scripts. The preprocessed MEG responses were then downsampled to 200 Hz before further analysis. The data were bandpass filtered between 1 and 30 Hz offline. An independent component analysis (ICA) was performed to remove artifacts including cardiac, eye movements, blinks, and environmental noise.

The continuously recorded MEG data were divided into epochs of 3 sec length (1 sec before and 2 sec after the cue word presentation) for the reactivation condition and 5 sec length (1 sec before and 4 sec after the substitute word presentation) for the interference condition. Baseline correction was applied by subtracting the average response of the 1 sec prior to the word presentation from all data points throughout the epoch.

Spectral analysis was performed on single trials and then averaged across trials. We used wavelet toolbox functions to examine the spectrotemporal power profiles. Specifically, the time-frequency representations (TFRs) was transformed using the continuous complex Gaussian wavelet (order = 4; e.g., FWHM = 1.32 s for 1 Hz wavelet) transform (Wavelet toolbox, MATLAB), with frequencies ranging from 1 to 30 Hz in increments of 1 Hz, for each condition, on each channel, and in each subject respectively. TFRs were then log10 transformed. Baseline correction for each frequency was applied by subtracting the averaged absolute values of the 0.5 sec prior to the word presentation from all data points throughout the epoch. Further analysis was performed only on planar gradiometers, considering that planar gradient maxima are located above neural sources which facilitates the interpretation of MEG results (Bastiaansen & Knösche, 2000; Hari & Salmelin, 1997). For spectral analysis, we computed metrics separately for the horizontal and vertical planar gradients, and combined the two by computing the sum.

Based on the previous findings in memory research (Klimesch, 1999), the theta- (5–7 Hz) and alpha-band (8–12 Hz) power profiles were extracted from the output of the wavelet transform for further analysis. For both reactivation (Fig. 2A) and interference (Fig. 3A) manipulations, six combined planar gradiometers showing the largest differences between the strong and baseline conditions at the theta and alpha bands were selected as sensor of interest for further analysis. Time of interest was determined by clustering the time points showing the most significant differences between the strong and baseline conditions at a step of 200 ms. An interval of 0.6 s was arbitrarily determined. The 1.0 −1.6 s after reactivation and 3.4 – 4.0 s after interference were selected. Correlations between neural signal change and behavioural effects were performed. To avoid the influence of outliers, we calculated the median absolute deviation (MAD) of each neural and behavioral response and excluded those that were outside three MADs. As a result, two participants with extreme alpha power changes during both reactivation and interference were excluded.

### Exps. 2–4

#### Participants and Materials

The sample sizes were based on our previous studies (see 30). Thirty-two (aged 19–31 years, 24 females), thirty-four (aged 19–28 years, 21 females) and thirty-two participants (aged 19–28 years, 21 females) participated Exps. 2, 3 and 4, respectively. They were all native Chinese speakers, with normal reading and comprehension ability, and no known neurological disorders. Participants gave written, informed consent in accordance with the procedures and protocols approved by the human participant review committee of Peking University.

Materials were 60 Chinese word pairs. The double-cue/one-target paradigm was used. Therefore, the 60 word pairs were composed by 60 cues and 30 targets, each target being paired with two different cues. All the words were frequency-balanced two-character Chinese words.

#### Design

The experiment consisted of three phases: associative learning, forgetting training, and recall testing. Both retrieval suppression and R-interference were used during forgetting training. To test the possible involvement of inhibition in R-interference, the memory impairments by suppression and interference were compared within participants.

#### Procedure

##### Exp. 2

###### Associative Learning

Participants learned 60 word pairs (30 A-X and 30 B-X pairs). Each word pair was first presented on the screen for 3 s, interleaved by a fixation cross for 1 s. To assist learning, a test-feedback session was given afterwards, which repeated until participants could recall all the target words when given the cue words.

###### Forgetting Training

Training was given seven days later, on A-X pairs. The original memory was first reactivated by presenting all A-X pairs to participants sequentially, each for 2 s. The 30 A-X pairs were then divided into three groups, 10 pairs each. One group accepted interference training, during which each cue A word was shown along with a substitute word. Participants were asked to memorize the new word pairs. One group was used for retrieval suppression, during which the cue words were presented alone on the screen and ‘no-think’ instructions were given. Specifically, participants were instructed to focus on the cues while suppressing the retrieval of the target items. The remaining group was not shown during this phase, and served as the baseline.

Reactivation was done once, and interference/suppression training repeated 12 times. The order for R-interference and R-suppression was counterbalanced within participants. The three subsets of A-X pairs were counterbalanced between participants for different conditions.

###### Recall Testing

A recall test was given five minutes later, on both the A-X and B-X pairs. Each cue word was presented on the screen sequentially. Participants were asked to retrieve the target words that were originally associated with the cues. Words from different conditions were presented in a pseudorandom order. No time limit was given.

##### Exp. 3

The procedure was the same as that in Exp. 2, except that participants reported the intrusion of the original memory right after each interference/suppression trial.

##### Exp. 4

The procedure was the same as that in Exp. 3, except that no pre-training reactivation was given. Namely, participants accepted interference and suppression training on two groups of A-X pairs directly.

**Fig. S1.**
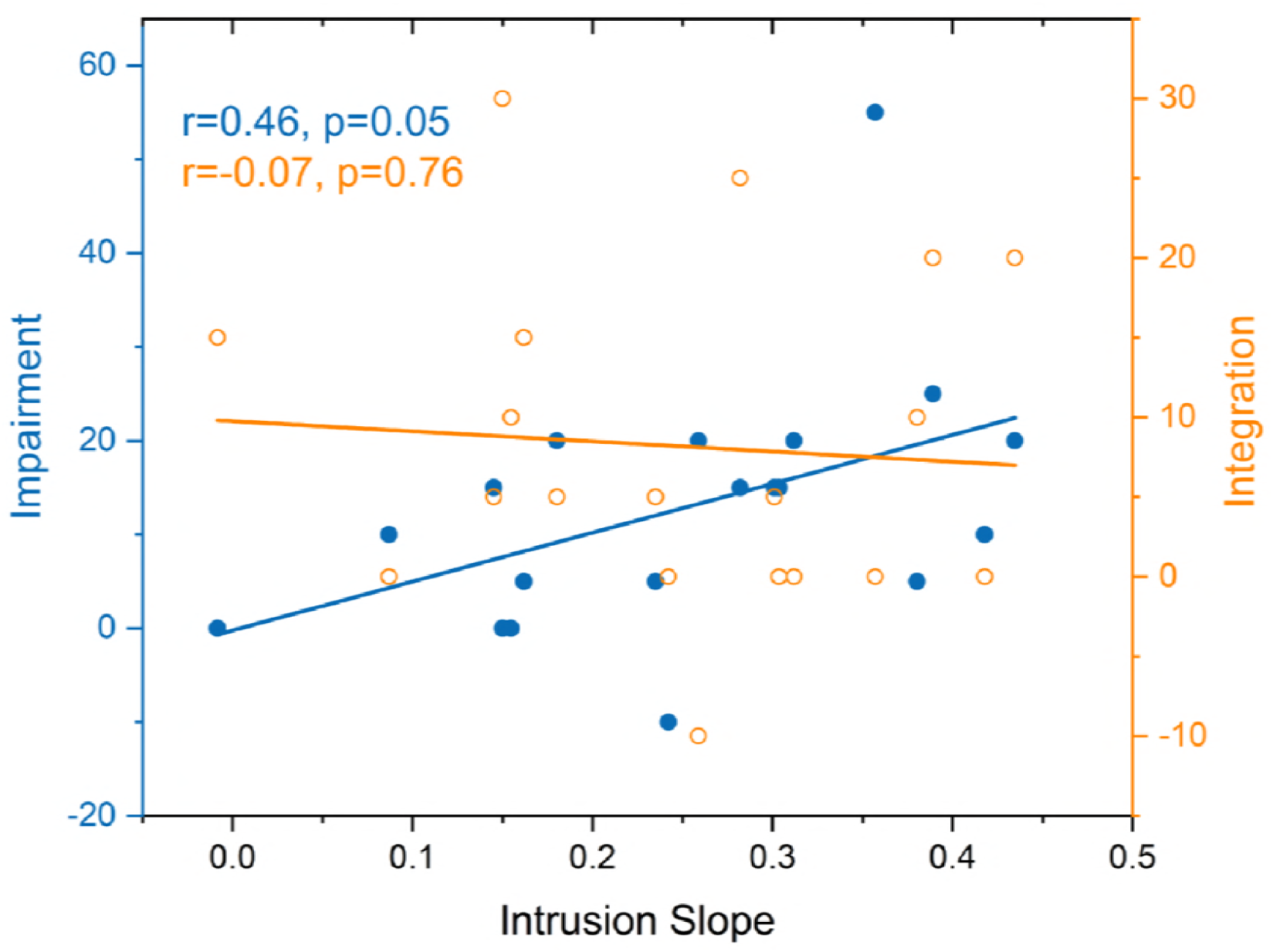
Correlation between memory impairment and integration with the slope of intrusion decrease in Exp. 1.

**Fig. S2.**
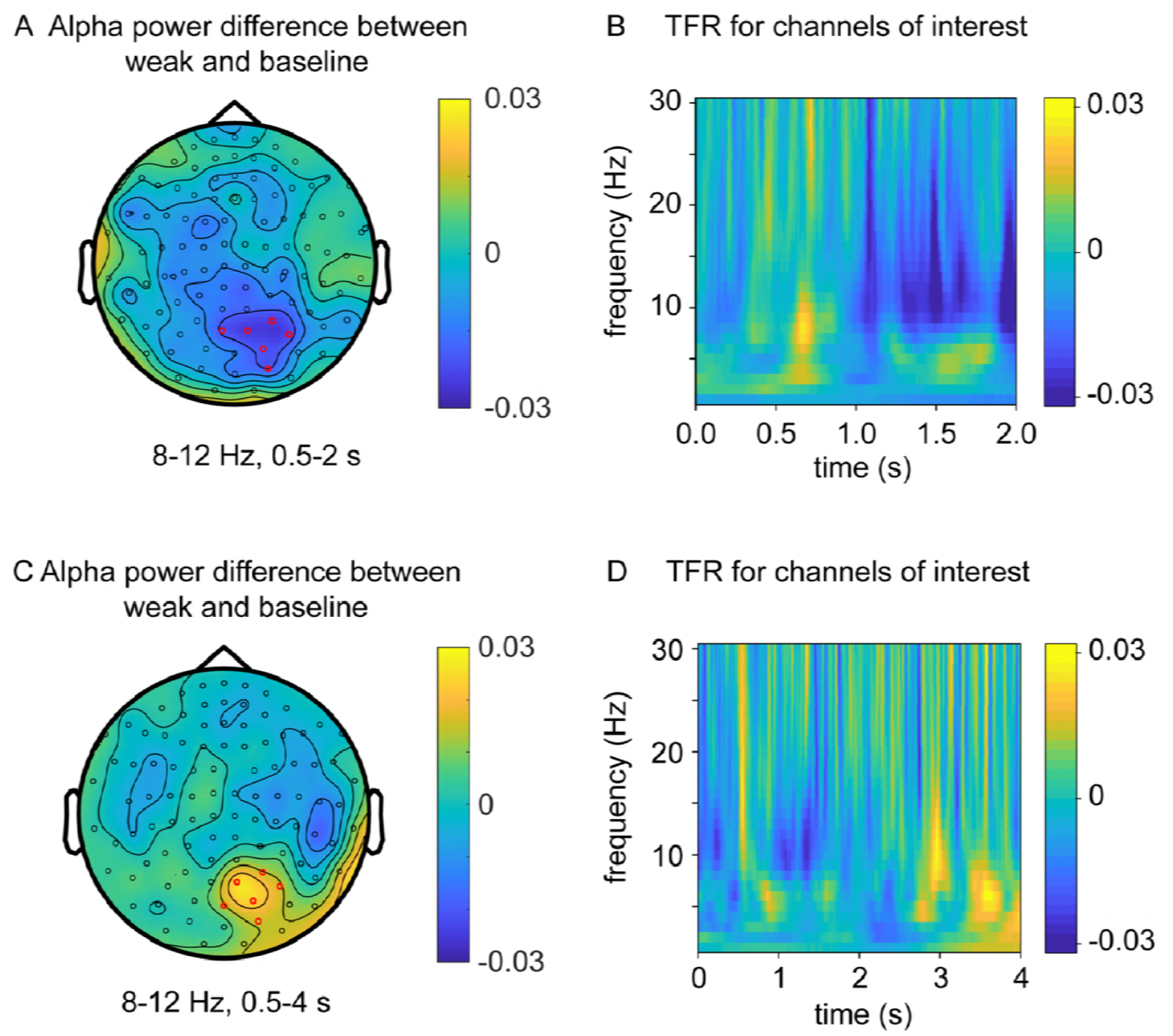
MEG results for the weak reactivation condition. (A) Grand average (N = 19) topographic map for alpha-band inhibition during reactivation (mean alpha-band power difference between weak and baseline conditions within 0.5 - 2 s). Red circles indicate channels with the strongest alpha decrease. (B) Grand average TFR profile of the selected six channels for the difference between weak and baseline conditions as a function of time (0 - 2s) and frequency (0–30 Hz). Alpha power decreased within the last 1 s of the reactivation phase. (C) Grand average (N = 19) topographic map for alpha-band enhancement during interference (mean alpha-band power difference between weak and baseline conditions within 0.5 - 4 s). Red circles indicate channels with the strongest alpha decrease. (D) Grand average TFR profile of the selected six channels for the difference between weak and baseline conditions as a function of time (0 - 4s) and frequency (0–30 Hz). Alpha power increased during the last 1 s of the interference phase.

**Fig. S3.**
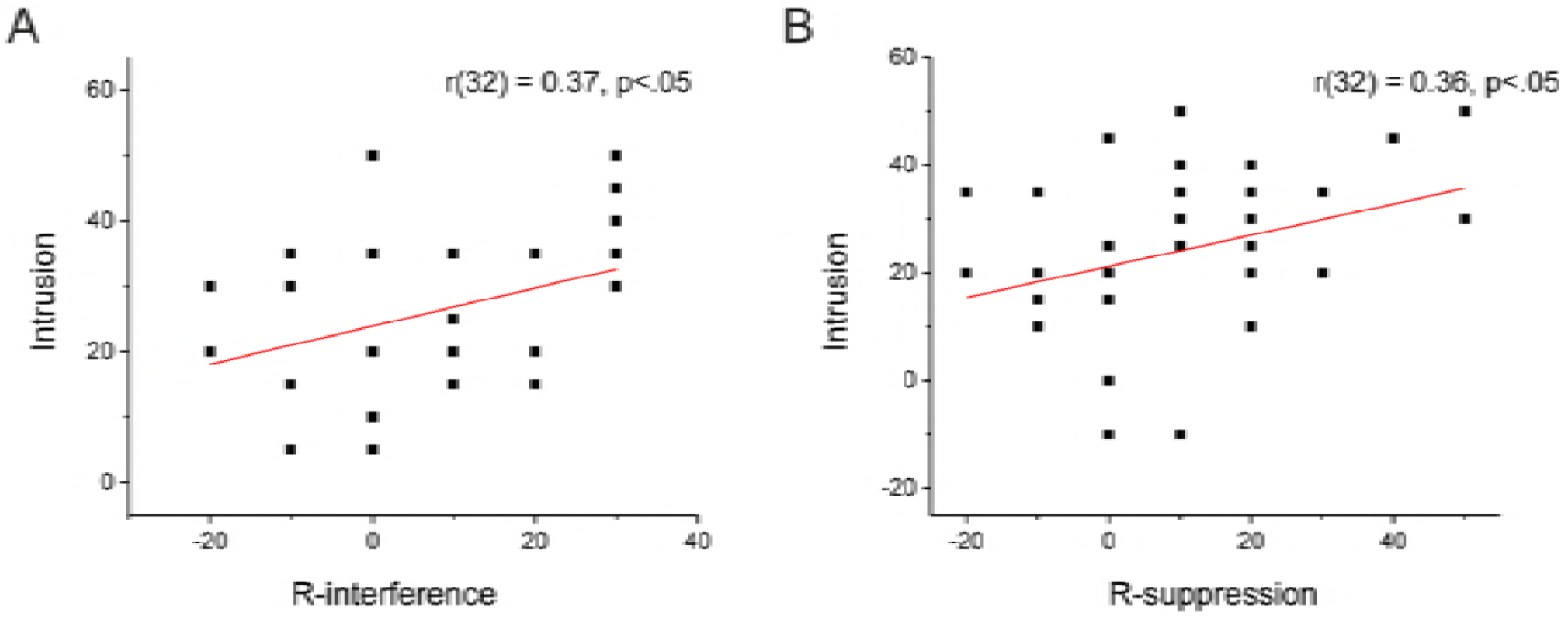
Correlations between the original memory intrusion and the impairment by R-interference (A) and R-suppression (B).

**Fig. S4.**
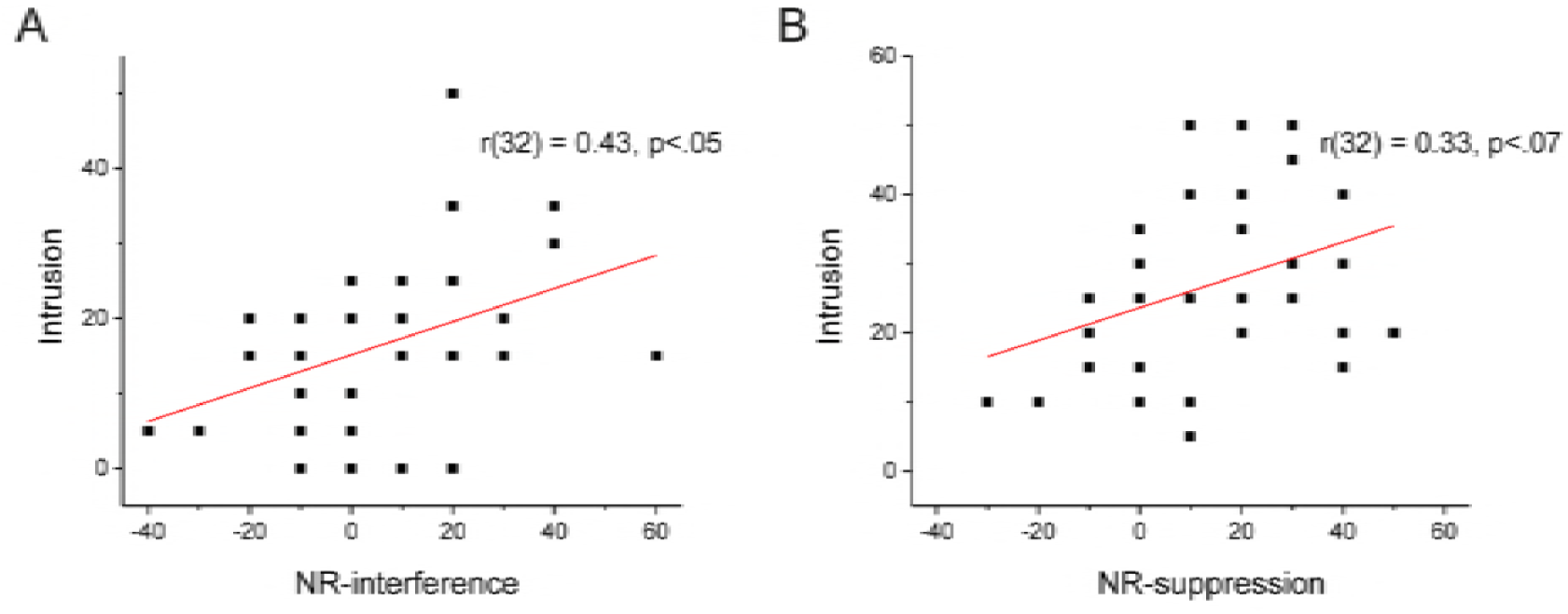
Correlations between the original memory intrusion and the impairment by NR-interference (A) and NR-suppression (B).

## Acknowledgements

This work was supported by National Natural Science Foundation of China (31421003, 31771205, 61690205), National Program on Key Basic Research Project (973 Program, 973-2015CB351800), the Peking-Tsinghua Center for Life Sciences, and Beijing Advanced Innovation Center for Genomics at Peking University.

